# A Bayesian Framework for Genome-wide Circadian Rhythmicity Biomarker Detection

**DOI:** 10.1101/2024.10.28.620703

**Authors:** Haocheng Ding, Lingsong Meng, Yutao Zhang, Andrew J. Bryant, Chengguo Xing, Karyn A. Esser, Li Chen, Zhiguang Huo

## Abstract

Circadian rhythms are endogenous ∼24-hour cycles that significantly influence physiological and behavioral processes. These rhythms are governed by a transcriptional-translational feedback loop of core circadian genes and are essential for maintaining overall health. The study of circadian rhythms has expanded into various omics datasets, necessitating accurate analytical methodology for circadian biomarker detection. Here, we introduce a novel Bayesian framework for the genome-wide detection of circadian rhythms that is capable of incorporating prior biological knowledge and adjusting for multiple testing issue via a false discovery rate approach. Our framework leverages a Bayesian hierarchical model and employs a reverse jump Markov chain Monte Carlo (rjMCMC) technique for model selection. Through extensive simulations, our method, BayesCircRhy, demonstrated superior false discovery rate control over competing methods, robustness against heavier-tailed error distributions, and better performance compared to existing approaches. The method’s efficacy was further validated in two RNA-Sequencing data, including a human resitrcted feeding data and a mouse aging data, where it successfully identified known and novel circadian genes. R package “BayesianCircadian” for the method is publicly available on GitHub https://github.com/jxncdhc/BayesianCircadian.

## Introduction

Circadian rhythms, endogenous cycles lasting ∼24 hours, evolved as adaptations to the natural light-dark cycles resulting from the Earth’s rotation. They regulate essential functions of physiological and behavioral processes such as sleep-wake patterns, body temperature fluctuations, and the secretion of melatonin [1, 19, 2, 7], impacting overall health and susceptibility to diseases. Understanding circadian rhythms, is crucial for comprehending human health and well-being. Circadian regulation lies a clock mechanism present in nearly all cells of the body. This mechanism operates through a transcriptional-translational feedback loop governed by a set of core clock genes [18, 13], which include *CLOCK, BMAL1* as transcriptomic activator, and the period family (*PER1, PER2, PER3*), as well as the cryptochrome family (*CRY1, CRY2*) as the inhibitors. The circadian transcriptomic landscape has been extensively explored across various tissues, including the postmortem brain [3, 33], skeletal muscle [14], liver [16], and blood [29]. Studies employing pan-tissue transcriptomic analyses in humans [32], mice [40], and baboons [30] have unveiled tissue-specific circadian patterns in gene expression. Furthermore, beyond transcriptomics, circadian investigations have extended to other omics data types, including DNA methylation [22], ChIP-Seq [20], proteomics [36], and metabolomics [5]. Accurate detection and analysis of circadian rhythms are imperative for numerous biomedical applications, including chronotherapy, personalized medicine, and the management of sleep disorders [28].

According to the literature, several algorithms have been developed to identify circadian rhythmicity. The cosinor-based model, for instance, proposes that the expression level of a gene follows a sine or cosine function of circadian time [4]. This model offers biological interpretability and accurate statistical inferences under Gaussian assumptions [8]. [27] introduced a cosinor mixed model for circadian rhythm data [27]. Other parametric approaches, such as Lomb-Scargle periodograms [11] and COSOPT [34], detect oscillating genes with irregular shapes by assuming mixtures of sine/cosine curves with distinct frequencies. In contrast, nonparametric models like ARSER [39], RAIN [35], and JTK CYCLE [17] are devoid of modeling assumptions, making them adept at capturing irregular curve shapes. MetaCycle [38] combines results from ARSER, JTK CYCLE, and Lomb-Scargle using meta-analysis via Fisher’s method. Both parametric and nonparametric algorithms have been widely used in circadian transcriptomic studies, and there several review studies systematically compare their performance [15, 25, 21].

The surge in the popularity of circadian omics approach has led to a proliferation of studies and consequent data deposit in public repositories. This accumulation of knowledge presents an exceptional opportunity to enhance our understanding the role of the molecular clock underlying various diseases and physiological processes. Although numerous algorithms for detecting circadian rhythmicity are available, there is a research gap: the majority adhere to a frequentist approach, neglecting the integration of established circadian knowledge. Consequently, there’s an significant need for a novel Bayesian approach that can incorporate this prior knowledge, which will enhance statistical power and reliability for detecting circadian biomarker in studies with similar experimental design. In addition, the Bayesian method also offers a framework to estimate uncertainty in predictions or parameter evaluations, which can be invaluable in understanding the variability and robustness of circadian biomarkers. Furthermore, it allows more flexible model specification compared to frequentist methods by enabling the use of hierarchical models to account for varying levels of data hierarchy, making them a promising tool for handling complex designs in circadian transcriptomic research.

To close the research gap, we developed a novel Bayesian framework for genome-wide circadian rhythm detection. Our framework is capable of incorporating prior knowledge, such as the identification of circadian genes from previous studies, into the modeling framework. By borrowing this information, we can greatly enhance the likelihood of accurately selecting a true circadian gene. To facilitate model selection between the circadian model and the non-circadian model, we adopted the reverse jump Markov chain Monte Carlo (rjMCMC) model, which is an extension special case of the Metropolis-Hastings algorithm when the dimension of model’s parameter space is not fixed [6]. This method allows the selection between two proposed circadian model and non-circadian model which have different dimension of parameters.

We evaluate our method under comprehensive simulation settings and compare it with other existing methods. We further apply our method on two real data applications including a time-restricted feeding data (RNA sequencing data of human skeletal muscles), and a lung mouse aging data (RNA sequencing data of the lung tissue of mice) The contribution and novelty of this paper includes: (i) First Bayesian method in circadian rhythmicity detection with improved accuracy with using of prior information; (ii) systematically compared the accuracy of the false discovery rate (FDR) in detecting circadian rhythmicity of our Bayesian detection method with other existing methods; (iii) our developed R package ‘BayesianCircadian’ is publicly available on GitHub, to promote its applications in the circadian field.

## Method

We develop a Bayesian Framework for detecting circadian biomarker for genome-wide transcriptomic data. To correct for the multiple testing, the decision framework (i.e., determining if a gene is a circadian gene) was developed based on false discovery rate. In addition, the method also allows to incorporate available prior information to enhance the detection of circadian genes.

### Notations for a cosinusoidal wave fitting

Our model assumes that the relationship between the gene expression value and the circadian time follows a cosinusoidal wave curve. As illustrated in Figure 1, denote *y* as the expression value for a gene; *t* as the circadian time; *A* as the amplitude; *M* as the MESOR (mid-line estimate statistic of rhythm). *ω* is the frequency of the Cosinusoidal wave, where 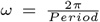. Without loss of generality, we set *period* = 24 hours to mimic the diurnal period. *ϕ* is the phase (i.e., peak time *t*_*p*_) of the cosinusoidal wave curve. Whenever there is no ambiguity, we will omit the unit “hours” in period, phase, and other related quantities.

**Fig 1.**
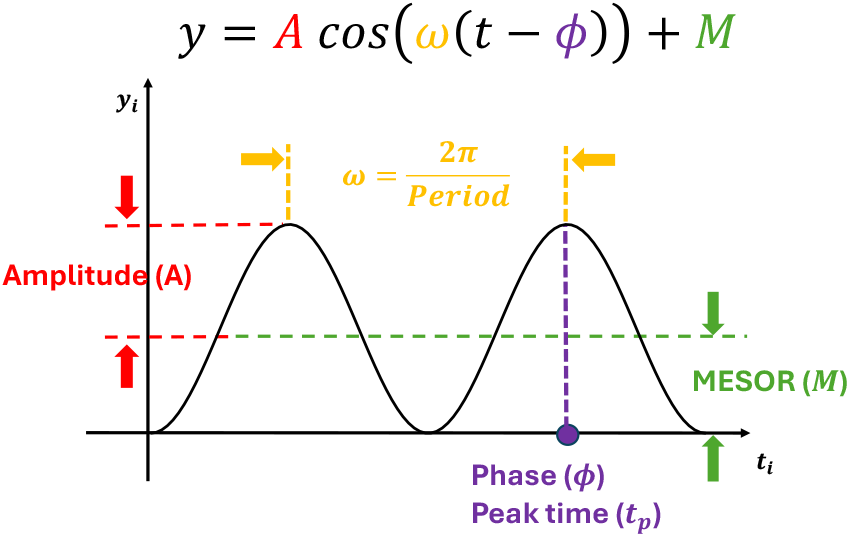
Illustration of a Cosine wave fitting and explanation of terminologies.

Denote *y*_*i*_ is the expression value of one gene for subject *i*(1 ≤ *i* ≤ *n*), where *n* is the total number of subjects. *t*_*i*_ is the circadian time for subject *i*. Under the circadian model *m*_1_, we assume the expression value of the circadian gene is:

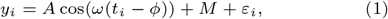

where *ε*_*i*_ is the error term for subject *i*; we assume *ε*_*i*_’s are identically and independently distributed (i.e., *iid*) and *ε*_*i*_ ∼ *N*(0, *σ*^2^), where *σ* is the noise level.

And under the non-circadian model *m*_0_, the expression value of the non-circadian gene is:

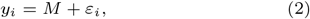

For the ease of discussion, we re-write the Equation 1 as:

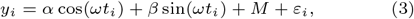

where *α* = *A* cos(*ωϕ*), and *β* = *A* sin(*ωϕ*).

### Bayesian hierarchical model

Leveraging the definition of *m*_0_ and *m*_1_, we further assume a Gaussian mixture model for gene *g*

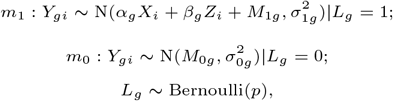

where *L*_*g*_ is a model indicator for each gene *g, L*_*g*_ = 1 indicates that the gene *g* is a circadian gene, *L*_*g*_ = 0 indicates that the gene *g* is a non-circadian gene; *Y*_*gi*_ denotes the expression value for each gene *g* and sample *i*; *X*_*i*_ = cos(*ωt*_*i*_) is the cosine transformation of circadian timepoint for each gene *g* and sample *i*; *Z*_*i*_ = sin(*ωt*_*i*_) is the sine transformation of circadian time for each gene *g* and sample *i*; *α*_*g*_ and *β*_*g*_ are the coefficients of *X*_*i*_ and *Z*_*i*_ respectively when gene *g* is a circadian gene; *M*_0*g*_ and *M*_1*g*_ are the MESOR for model *m*_0_ and *m*_1_ respectively. 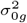 and 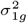 are the variance for model *m*_0_ and *m*_1_ respectively.

Figure 2 shows the graphical model representation of the data generative process of our Bayesian hierarchical model. Denote the parameter space for *m*_0_ as 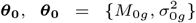. Denote the parameter space for *m*_1_ as ***θ***_**1**_, 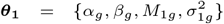. The parameters of the entire Bayesian hierarchical model **Θ** = {***θ***_**0**_, ***θ***_**1**_, *p, L*_*g*_} includes *α*_*g*_, *β*_*g*_, *M*_1*g*_, *M*_0*g*_, 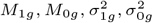, *p, L*_*g*_.

**Fig 2.**
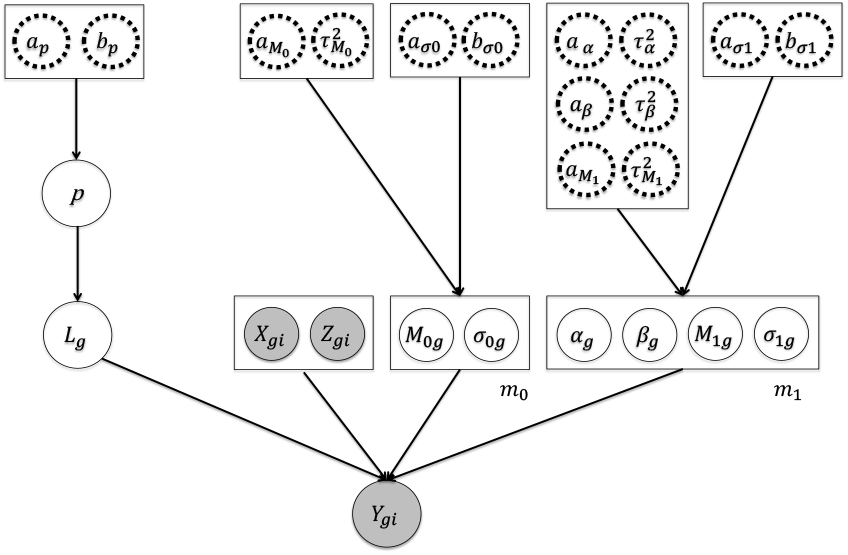
Graphical representation of Bayesian latent hierarchical model. Shaded nodes indicate observed variables. Dashed nodes indicate pre-fixed hyper parameters. Arrows show generative process. *g*(1 *≤ g ≤ G*) is the gene index, *i*(1 *≤ i ≤ n*) is the sample index.

### Prior specification

We apply independent conjugate priors to each component if a gene is a circadian gene) was developed based on false discovery rate. In addition, the method also allows to incorporate available prior information to enhance the detection of circadian genes. in **Θ** as follows: 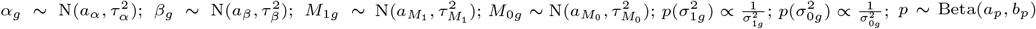. Such design of conjugate priors will greatly facilitate efficient implementations of Gibbs sampling. Here, 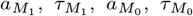, *a*_*α*_, *τ*_*α*_, *a*_*β*_, *τ*_*β*_, 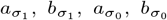, *a*_*p*_, *b*_*p*_ are hyper parameters.

### Hyper parameter justification

Where there is no prior data information, we propose to assign non-informative hyper parameters wherever possible. To be specific, we set *a*_*α*_ = 0, *τ*_*α*_ = 10; *a*_*β*_ = 0, *τ*_*β*_ = 10; 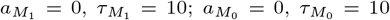; *a*_*p*_ = *b*_*p*_ = 1. Such non-informative hyper parameters will facilitate a data-driven approach.

### Reverse jump Markov chain Monte Carlo

Since the number of parameters are different in *m*_0_ and *m*_1_, and a gene can only come from one of the two models, it is challenging to directly use the conventional Bayesian sampling techniques to obtain the distribution of the Bayesian hierarchical model because of the transdimensional issue. To solve this challenge, we will leverage the reverse jump Markov Chain Monte Carlo (rjMCMC). As introduced by [12], reverse jump Markov Chain Monte Carlo (rjMCMC) is an extension of the traditional Metropolis-Hastings (MH) algorithm [26]. It’s suitable when exploring models of different complexities, allowing to compare models with different numbers of parameters or different structures. In particular, to solve the transdimensional issue, we introduce auxiliary variables 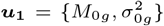 and 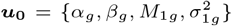 such that (***θ***_**0**_, ***u***_**0**_) = (***u***_**1**_, ***θ***_**1**_). In standard MH algorithm, we draw new samples from a proposal distribution, and decide whether to accept or reject the new samples based on certain acceptance ratio. In rjMCMC, in addition to draw samples, we also draw new models and the acceptance ratio is determined by both the samples and models. To be specific, the acceptance ratio for rjMCMC for a transition from *m*_*k*_ to 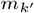 is:

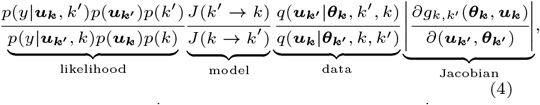

Where *k* and *k*^′^ are the model index with *k, k*^′^ ∈ {0, 1}. 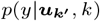 represents the likelihood of model *m*_*k*_, *p*(***u***_***k***_) represents the prior probability of the parameters of model *m*_*k*_, And *p*(*k*) represents the prior probability of model *m*_*k*_.

*J*(*k*^′^ → *k*) is the transitional probability of the model *m*_*k*_ from 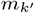. 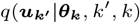, *k*^′^, *k*) is the data proposal density using model *m*_*k*_ [23]. 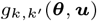 (***θ, u***) is the variable mapping function between model *k* and *k*^′^ (i.e., 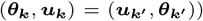, thus the Jacobian term equals to 1. Note that when there is no model jump (i.e., *k* = *k*^′^), it can be derived that this acceptance ratio is always 1, thus the algorithm reduces to a conventional Gibbs sampling.

And the implementation of this rjMCMC has been proposed by [10] as the following:

1. Initialize *L*_*g*_ and 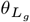 at iteration *j* = 1.
2. For iteration *j*(*j >* 1) perform:
  a. Temporary within-model move: for both *m*_1_ and *m*_0_, update the parameters ***θ***_**1**_ and ***θ***_**0**_ according to certain Gibbs sampler updating scheme.
  b. Temporary between-models move: simultaneously update model indicator *L*_*g*_ and the parameters 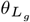 according to specific reversible proposal/acceptance mechanism.
  c. Increment iteration *j* = *j* + 1. If *j < N*_*MCMC*_, where *N*_*MCMC*_ is the total number of iterations, repeat step 2.

### Posterior calculation

Assume the state of the Markov chain for gene *g* is {*L*_*g*_(*j*), ***θ***_*Lg*_(*j*)} at current iteration *j*, then the update through rjMCMC algorithm at next iteration *j* + 1, where we also use and ***u***_**0**_ to indicate the temporary coefficient 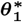

### Temporary within-model move

For circadian model *m*_1_, we calculate the temporary coefficient 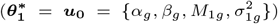, where we use ***u***_**0**_ and 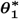 interchangeably to indicate the temporary coefficient.

1. Update temporary coefficient *α*_*g*_ from

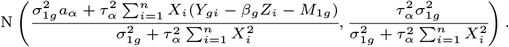
2. Update coefficient *β*_*g*_ from

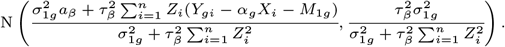
3. Update intercept *M*_0*g*_ from

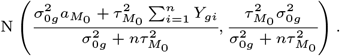
4. Update variance 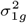 for circadian genes from

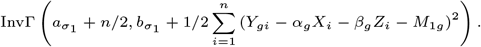

Similarly, for non-circadian model *m*_0_ we calculate the temporary coefficient 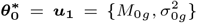, where we also use ***u***_**1**_ and 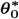 interchangeably.

1. Update intercept *M*_0*g*_ from

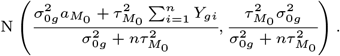
2. Update variance 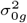
for non-circadian genes from

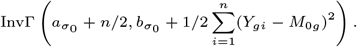

### Temporary between-model move

1. Update the proportion of circadian genes *p* from

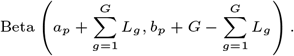
2. Update temporary model indicator 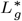 from the Bernoulli distribution with probability *prob*, i.e., Bern(*prob*), where

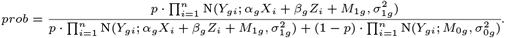
3. If 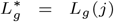, then gene *g* stays in model 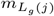 at iteration *j* + 1. There is no trans-model jump. The algorithm reduces to a typical Gibbs Sampler, and directly update the 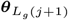 using the temporary values 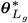 in the within-model move procedure for 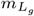.
4. If 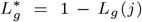, then gene *g* intends to jump from model 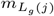 to model 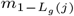 at iteration *j* + 1. In this case, we the acceptance ratio for this trans-model move is determined by the acceptance ratio detailed at the step 5 of section Temporary between-model move. If we accept, we simultaneously update *L*_*g*_(*j* + 1) = 1 − *L*_*g*_(*j*) and 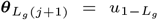, where 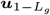 is determined in the temporary within-model move. If we don’t accept, the model and its parameters stay at *L*_*g*_(*j* + 1) = *L*_*g*_(*j*) and 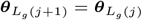.
5. By equation 4, acceptance probability of the new model can be calculated as the minimum between one and

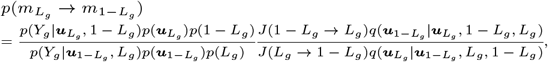

where 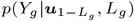 is the likelihood at iteration 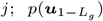 is the prior for parameters at iteration *j*; *p*(*L*_*g*_) is the prior for circadian gene proportion at iteration *j*; *J*(*L*_*g*_ → 1 − *L*_*g*_) is the model proposal probabilities at iteration 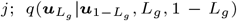 is the proposal densities for auxiliary variables at iteration *j*.

For example, considering the case that *L*_*g*_ = 0 at iteration *j* and *L*_*g*_ = 1 at iteration *j* + 1, it involves model transition from *m*_0_ to *m*_1_. Here are the steps to calculate the acceptance probability *p*(*m*_0_ → *m*_1_):

a. 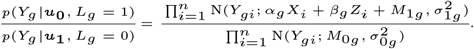
b. 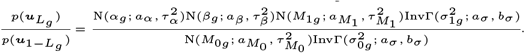
c. 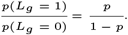
d. 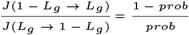

where *prob* was defined in section Temporary between-model move Step 2.
e. 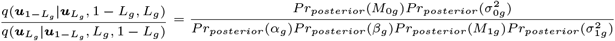

where

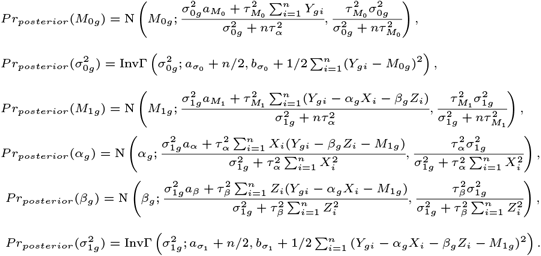

### Decision framework

We use the false discovery rate (FDR) approach to construct the decision rule to declare whether a gene is a circadian gene or not. To infer if a gene is a circadian gene, we first denote Ω_*C*_ as the collection of circadian genes (i.e., Ω_*C*_ = {*g* : 1 ≤ *g* ≤ *G*}; gene *g* is a circadian gene), and Ω_*N*_ as the collection of non-circadian genes (i.e., Ω_*N*_ = {*g* : 1 ≤ *g* ≤ *G*; *g* ∈*/* Ω_*C*_}). We denote *P*_*g*_ = Pr(*g* ∈ Ω_*N*_ |*L*_*g*_ = 1), which is also referred as the local discovery rate [9]. Given a threshold *η* (0 *< η <* 1), when declaring gene *g* as a circadian gene if *P*_*g*_ ≤ *η*, the expected number of false discoveries is Σ _*g*_ *P*_*g*_*I*(*P*_*g*_ ≤ *η*). According to [31], the expected false discovery rate for genes with *P*_*g*_ ≤ *η* is:

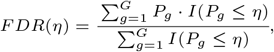

In practice, *P*_*g*_ is estimated as 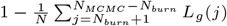, where *N*_*burn*_ is the number of burn-in samples, *N*_*MCMC*_ is the total number of rjMCMC iterations and *j* indicates the index of rjMCMC iteration.

### Incorporate prior knowledge

Our model also allows the incorporation of prior information of a gene being circadian or not. In our Bayesian hierarchical modeling framework, *p* represents the proportion of circadian genes of the entire genome. When we have prior knowledge, assume Ψ_*C*_ as the prior collection of circadian genes (i.e., Ψ_*C*_ = {*g* : 1 ≤ *g* ≤ *G*; gene *g* is a circadian gene from the prior knowledge}), and Ψ_*N*_ as the prior collection of non-circadian genes (i.e., Ψ_*N*_ = {*g* : 1 ≤ *g* ≤ *G*; *g* ∈*/* Ψ_*C*_}). Consequently, |Ψ_*C*_ | = *G* − |Ψ_*N*_ |, where *G* is total number of genes, |Ψ_*C*_ | and |Ψ_*N*_ | are the numbers of genes in Ψ_*C*_ and Ψ_*N*_, respectively.

Instead of assuming a uniform *p* across the genome, we propose *p*_*C*_ as the proportion of circadian genes within Ψ_*C*_ (i.e., *g* ∈ Ψ_*C*_), and *p*_*N*_ as the proportion within Ψ_*N*_ (i.e., *g* ∈ Ψ_*N*_). Under this framework, genes from the prior circadian gene list Ψ_*C*_ are better poised to borrow information from other genes in the same category, enhancing their likelihood of being classified as circadian genes. For example, if in a prior study, Ψ_*C*_ is the putative circadian gene set under false discovery rate threshold *p*_*F DR*_, then we could assume the prior for *p*_*C*_ is from 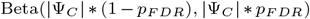. The expected values for *p*_*C*_ would then be 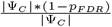, aligning with the observed proportion of true circadian genes. Similarly, we assume *p*_*N*_ follows an non-informative Beta(1, 1) prior distribution with mean 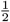.

### Other competing methods

We will compare our proposed Bayesian method for circadian biomarker detection to other existing methods, including, ARSER [39], Lomb-Scargle periodograms [11], JTK CYCLE [17], RAIN [35], MetaCycle [38], and LR rhythmicity [8]. Several of these methods have some special requirements for the input of circadian time: the input circadian time has to be integer value, and the intervals between two adjacent circadian time points must be equal. Thus, we will accommodate such design in our simulation settings when needed.

## Simulation

Based on the simulation result, we demonstrated that our proposed Bayesian method for circadian biomarker detection correctly controlled the false discovery rate to the 5% level, while some of the other methods failed to control the false discovery rate. In all later evaluations, we denoted “BayesCircRhy” as the Bayesian method for circadian biomarker detection, and “BayesCircRhy (Prior)” as the Bayesian method with incorporation of prior knowledge.

### Simulation to demonstrate the Bayesian method can control the FDR to the nominal level under various settings

#### Simulation settings

Denote *i*(1 ≤ *i* ≤ *n*) as the sample index, where *n* was the total number of samples. The circadian time *t*_*i*_ for sample *i* was generated from uniform distribution UNIF(0, 24). We simulated the gene expression value for sample *i* using Equation 1. Our basic parameter setting for simulation is listed as below. For each gene, the sample size *n* was 48; the circadian time were sampled from UNIF(0, 24). Amplitude *A* was set as a fix value at 3 for circadian gene and 0 for non-circadian gene; phase *ϕ* was generated from UNIF(0, 24). MESOR *M* was fixed at 3. Error term *ε*_*i*_ was generated from normal distribution *N* (0, *σ*^2^) where *σ*^2^ was set to be 1. We simulated *G* = 1, 000 genes with 100 circadian genes and performed 5,000 MCMC iterations for each simulation. Each simulation was repeated *B* = 10 times to increase the numbers of replications and obtain a standard error estimate. Additionally, we conducted following simulation settings to examine the robustness of our method against higher signal-to-noise ratios, variations in sample size, different proportions of circadian genes, and violations of normality assumptions.

1. Impact of sample size. We varied *n* = 24, 48, 96 while fixing other parameters in the basic parameter setting fixed.

2. Impact of signal noise ratio. The signal noise ratio is defined as 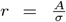, which can be viewed as the effect size of a circadian gene [41]. Thus we varied *r* = 2, 3, 4, 5 to mimic varying levels of signal noise ratio, while fixing other parameters in the basic parameter setting.

3. Impact of different proportions of circadian genes. In real data applications, datasets may have different percentages of circadian genes. Thus, we simulated the datasets contain percentage of circadian genes at 5%, 10%, 15% and 20%.

4. Violation of the Gaussian assumption. Instead of assuming the error term was generated from a standard normal distribution (i.e., *N* (0, 1)), we generated *ε*_1_ ∼ *t*(2), *t*(3), *t*(5), *t*(10), *t*(∞), where *t*(*df*) is the *t*-distribution with degree of freedom *df*. This family of *t*-distributions represents heavy-tailed error distribution, with smaller *df* indicating longer tailed error distribution, and thus larger violation of the normality assumption. When *df* = ∞, *t*(∞) becomes *N* (0, 1).

#### Simulation evaluations

We evaluated the performance of our proposed Bayesian circadian method (BayesCircRhy) in circadian rhythmicity detection while fixing the nominal FDR to be 5%.

As shown in figure 3a, while varying (i) number of sample size (*n*), BayesCircRhy controlled the FDR at the nominal 5% level for different sample sizes. In Figure 3b when varying (ii) signal noise ratio (*r*), BayesCircRhy successfully controlled the FDR at the 5% nominal level when *r* = 2 ∼ 5. (iii) with different number of circadian genes (*G*1) in figure 3c, BayesCircRhy controlled the FDR at 5% level for all settings; (iv) by varying the degree of freedom of the *t*−distributed errors (*t*(*df*)) in figure 3, we observed when there were mild (*t*(10)) and moderate (*t*(5)) violations of normality assumption, the FDR was in line with the 5% nominal level. However, when the violation of the normality assumption was severe (i.e., *t*(2)), the FDR was conservative (0.022).

**Fig 3.**
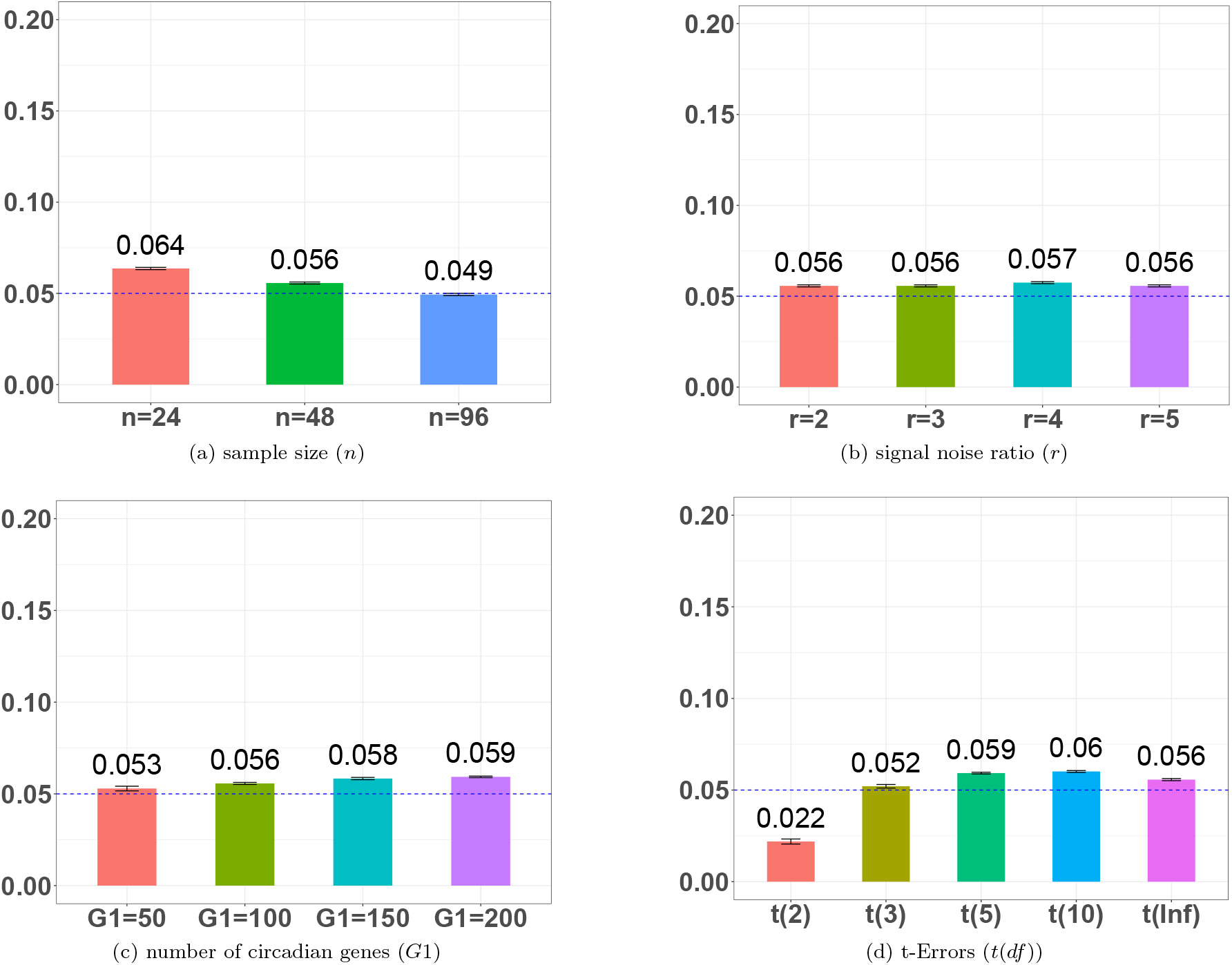
FDR evaluation for BayesCircRhy in detecting circadian rhythms. The simulations settings included varying sample size, level of noise ratio, number of circadian genes and *t*-distributed errors. The standard error of the mean FDR was also marked on the bar plot.

We further evaluated the genome-wide power of BayesCircRhy.

The genome-wide power is defined as 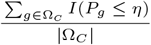, where Ω_*C*_ is the collection of circadian genes; 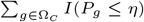 is the sum of circadian genes identified by BayesCircRhy and |Ω_*C*_ | is the total number of true circadian genes.

As shown in figure 4, BayesCircRhy demonstrated high power (93.2% − 100%) across all simulation settings.

**Fig 4.**
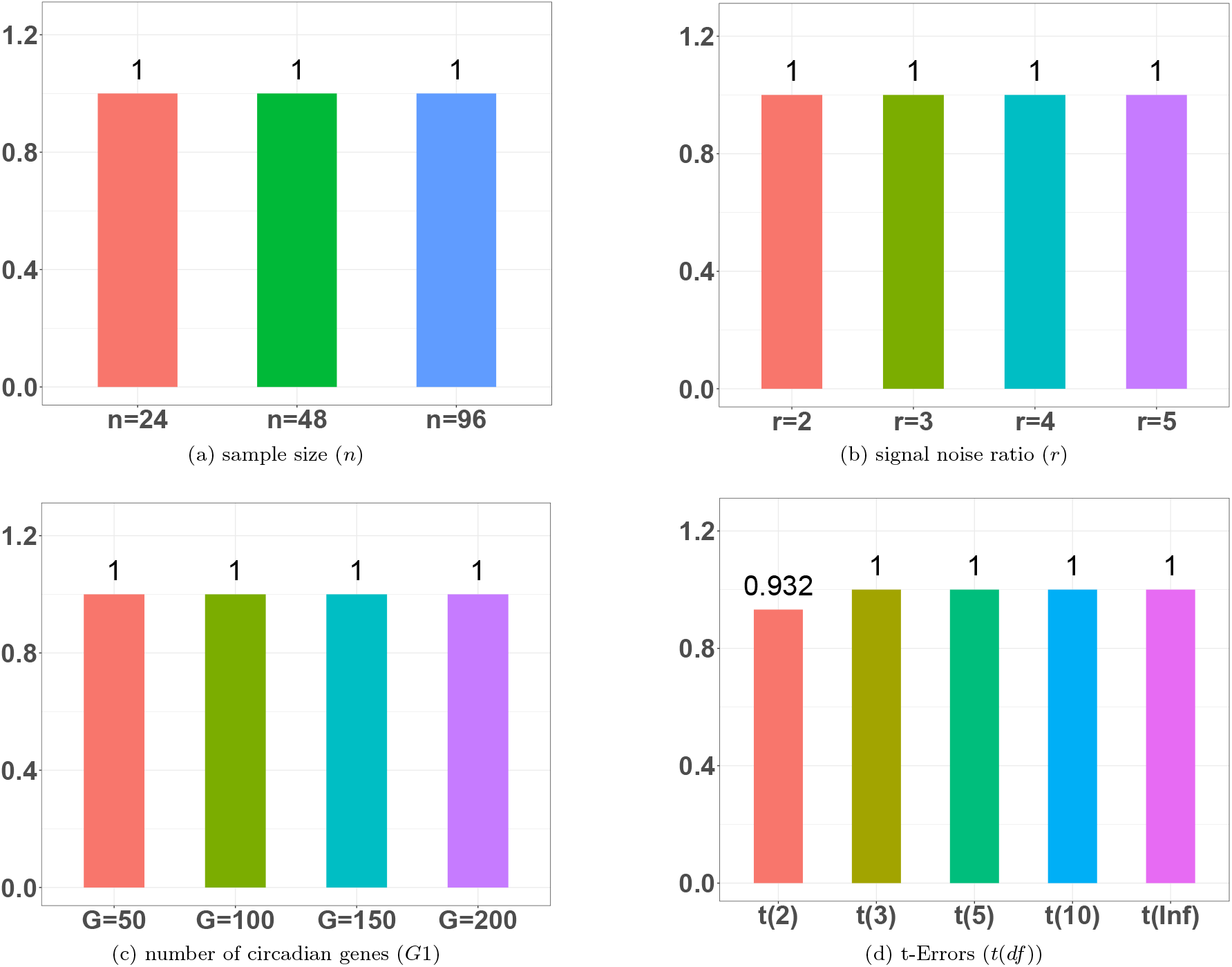
Power evaluation for BayesCircRhy in detecting circadian rhythms. The simulations settings included varying sample size, level of noise ratio, number of circadian genes and *t*-distributed errors.

### Simulation to compare with other competing methods

#### Simulation settings

To compare with other competing methods, we inherited the basic simulation section as detailed in section Simulation settings. We change the circadian time to be evenly sampled from 0 ∼ 24 (i.e., *t*_1_ = 1, *t*_2_ = 2, …, *t*_48_ = 24) which required by other existing methods.

We further included the result from BayesCircRhy incorporating prior information. To be specific, we generated two datasets with the basic simulation setting in section Simulation settings and labeled them as prior dataset and test dataset. First, we applied our Bayesian algorithm without priors to the test dataset. The result from this method is labeled as BayesCircRhy. Further, we generate prior knowledge by applying the Bayesian algorithm to the prior dataset, and incorporate its detection result as prior information to the test dataset. The result from this approach is labeled as BayesCircRhy (prior).

#### Simulation evaluations

We compared the BayesCircRhy and BayesCircRhy (prior) with other major existing methods including LR rhythmicity, Lomb-Scargle, JTK, ARSER, MetaCycle, and Rain in terms of detecting circadian rhythmicity. Figure 5 showed false discovery rates from different existing methods. In general, BayesCircRhy and BayesCircRhy with prior information controlled the false discovery rate to the 5% nominal level, while other methods had either inflated or conservative false discovery rate. Therefore, we will focus on our proposed Bayesian methods in the real data application.

**Fig 5.**
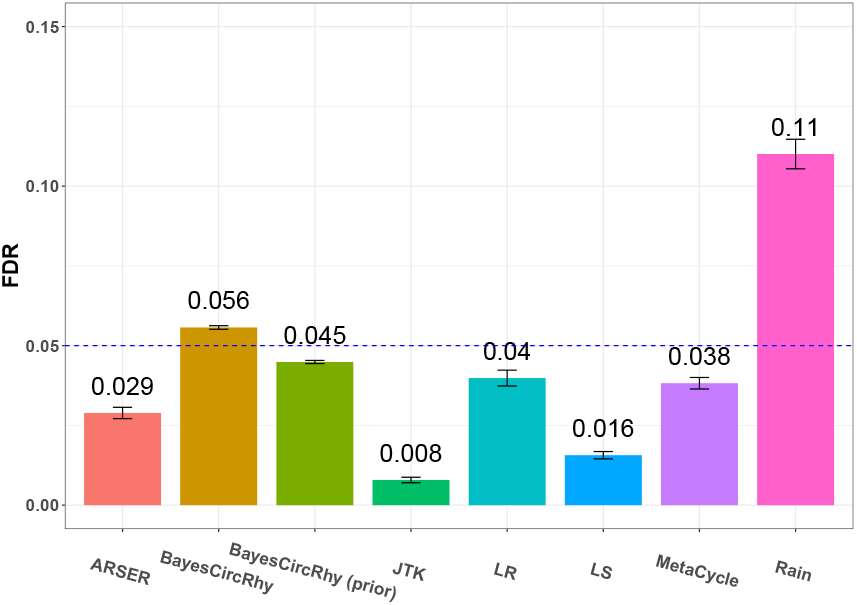
False discovery rate (FDR) comparison for 8 different methods in detecting circadian rhythmicity. The blue dashed line is the FDR 5% level. A higher than 5% blue dashed line bar indicates an inflated FDR; a lower than 5% blue dashed line bar indicates a conservative FDR; and a bar at the blue dashed line indicates a accurate FDR (i.e., *q* = 0.05). The standard error of the mean FDR was also marked on the bar plot. LR rhythmicity was denoted as LR. Lomb-Scargle was denoted as LS.

## Real data application

We evaluated the BayesCircRhy in two real data applications, including a human gene expression RNA sequencing data investigating time restricted feeding, and a mouse gene expression RNA sequencing data investigating aging. In this section, we used *q*-value *<* 0.05 as the cutoff to declare statistical significance unless otherwise specified. Since our proposed Bayesian approach performs better than the other competing methods in simulation, and in real data application, it is difficult to benchmark the performance of different methods, we opt out comparing with other existing methods.

### Human time-restricted feeding data

We evaluated the performance of our method in transcriptomic profiles of human skeletal muscle tissue. 11 overweight or obese male participants aged between 30 and 45, with body mass indices (BMIs) ranging from 27 to 35 *kg/m*^2^, were included in this study; Following a crossover design, participants were randomly allocated to either a time-restricted feeding (TRF) or an unrestricted feeding (URF) group, sequentially experiencing both regimens. The skeletal muscle samples of each participant under each experimental group were repeatedly measured every 4 hours over a 24-hour period. Each participant contributed 4 ∼ 6 measurements, yielding a total of 63 samples in the TRF group and 62 in the URF group. Detailed description of this study has been previously published [24]. The RNA-seq dataset generated from this study is publicly accessible through the Gene Expression Omnibus (GEO) database with accession number GSE129843. The gene expression data was normalized and log2 transformed. After filtering out lowly expressed genes, 13,169 genes were remained for further analysis.

Firstly, our analysis with BayesCircRhy focused solely on the samples from the URF group. By applying False discovery rate at 5%, we identified a total of 30 genes exhibiting significant circadian genes using BayesCircRhy. Figure 6 shows the top 6 significant circadian genes within the URF group, namely *PER1, PER2, PER3, ARNTL, NR1D1*, and *DBP*, which are well-recognized for their persistent circadian patterns [21]. All of them demonstrated significant *q*-values, showing the strong detection power of our method in identifying circadian rhythms.

**Fig 6.**
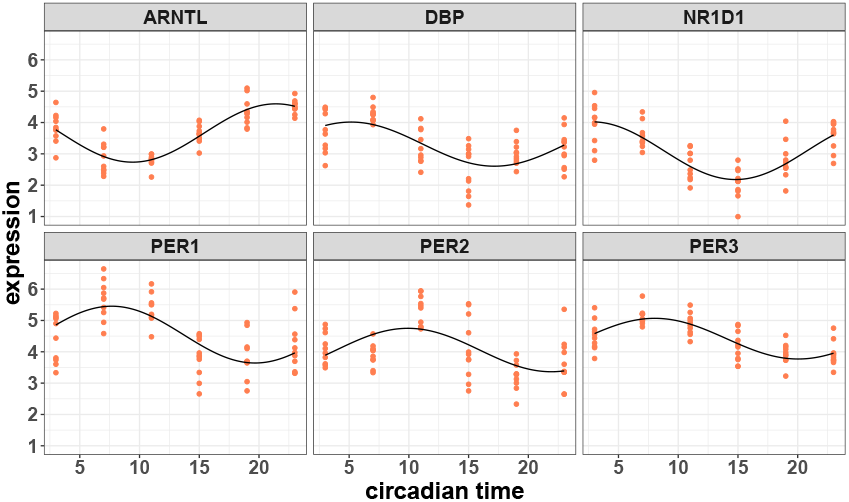
Circadian rhythmicity for top 6 significant circadian genes in the URF group of human restricted feeding data, including *ARNTL, DBP, NR1D1, PER1, PER2* and *PER3*, using BayesCircRhy.

To further examine the performance of BayesCircRhy with prior information, we utilized the TRF groups as the prior information. To be specific, we first apply the BayesCircRhy to the TRF groups, and identified 135 circadian genes at 5% FDR. Then we incorporated the putative circadian genes as the prior information for the BayesCircRhy method, and applied this method to the URF group. A Total of 75 circadian genes were identified, with 45 more genes comparing to the results with no such prior information. This is not unexpected since by borrowing information from the prior set, the BayesCircRhy is more likely to detect circadian genes.

### Mouse aging data

Additionally, we evaluated our method in a mouse aging data from lung tissue. The dataset profiled the circadian transcriptome in the lung tissue by 3 age groups young (6-month), aged (18-month) and old (27-month). All samples were male mice. Tissues were collected by every 4 hours for 48 hours in total darkness with the first time point being the circadian time 18 (CT18) and there were total of 12 time points. Detailed description of the dataset has been published [37]. The RNA-seq dataset generated from this study is publicly available on the Gene Expression Omnibus (GEO) database with accession number GSE201207. The data was normalized via count per million reads (CPM), followed by log2 transformation. We focused on the aged group with lung tissue using BayesCircRhy in our analysis. After filtering out lowly expressed genes, 15,896 gene probes were remained in the analysis.

At FDR 5%, we identified a total of 4 genes exhibiting significant circadian genes using BayesCircRhy in the aged group. To further examine the performance of BayesCircRhy with prior information, we utilized the young group as the prior information. To be specific, we first apply the BayesCircRhy to the young group in the lung tissue, and identified 15 circadian genes at 5% FDR. Then we incorporated the putative circadian genes as the prior information for the BayesCircRhy method, and applied this method to the aged lung group. A Total of 16 circadian genes were identified, with 12 more genes comparing to the results with no such prior information. Figure 7 shows the top 6 significant circadian genes within the aged group, namely *ADM, ARNTL, DBP, NR1D1, NR1D2* and *SPON2*.

**Fig 7.**
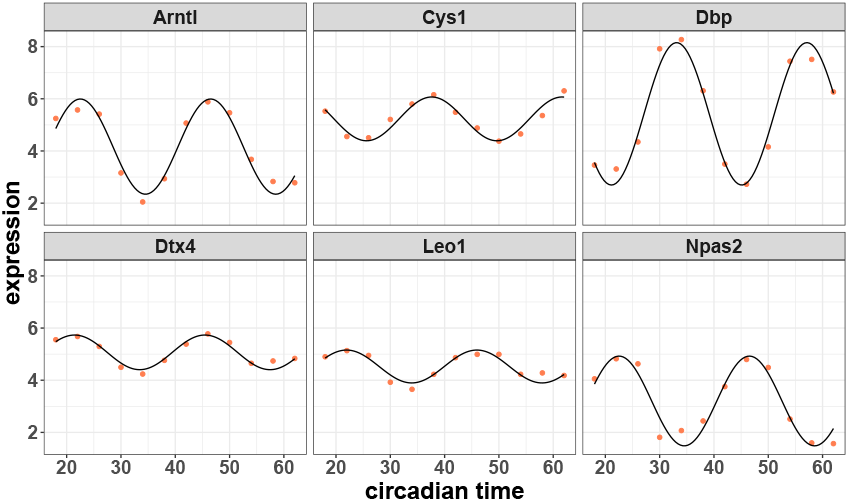
Circadian rhythmicity for 6 significant circadian genes in the lung tissue aged group of mouse tissue data, including *Arntl, Cys1, Dbp, Dtx4, Leo1* and *Naps2*, using BayesCircRhy.

## Discussion

In summary, we developed a Bayesian framework for circadian rhythmicity biomarker detection in genome-wide transcriptomic data (BayesCircRhy). The proposed Bayesian hierarchical model leveraged a mixture model approach to tackle the circadian model/null model selection problem. The transdimentional issue of the circadian model and the null model was solved using rjMCMC. Our full Bayesian approach allows efficient sampling of all model parameters, which enables the estimation of parameter uncertainties. We further employed the false discovery rate approach to facilitate the decision make process (to declare whether a gene is a circadian gene). In simulations, we demonstrated that the proposed method BayesCircRhy could control the FDR to the nominal 5% in most scenarios. And comparing with its competing methods, the BayesCircRhy has better performance in controlling the FDR to the nominal level. We further extended our Bayesian to incorporate prior biological knowledge. And in simulations and real data applications, we demonstrated that by incorporating prior knowledge, our approach can better capture the true circadian genes. We further demonstrated the superious performance of our method in transcriptomic data applications including two RNA-seq data, human time restricted feeding data and mouse multi-tissue data.

Our method has the following strengths. (i) When the prior data information is available, our method could incorporate the information to enhance the circadian biomarker detection process. (ii) BayesCircRhy is robust against the violation of the Gaussian assumptions. As shown in our simulation, even under moderate deviations from normality assumptions (i.e. *t*(3),

*t*(5) and *t*(10)), our method successfully maitained the FDR approximately 5%. (iii) Our method has been implemented in the R package ‘BayesianCircadian’, which is publicly accessible on GitHub (https://github.com/jxncdhc/BayesianCircadian).

Our method has the following limitations. (i) The Bayesian design assumes a cosinor based model and residuals are independently, and and normally distributed. Although in the simulation section, our method shows its robustness when there are some moderate violation of Gaussian assumption, sever deviations of these assumption may still result in an inflated/deflated FDR. Therefore, we will extend our framework toward the non-Gaussian error distribution, and repeated measurement observations as future work. (ii) Our approach is a little conservative when sample size or effect size is small. Nonetheless, our Bayesian framework is the first Bayesian framework for detecting circadian biomarkers in a genome-wide setting that can incorporate prior knowledge. This framework can be modified to adapt to many other complex circadian problems in transcriptomic studies.

## Author contributions statement

H.D. and L.M. contributed equally to the method development, data analysis, and R package preparation. H.D., Z.H. and

L.C. contributed to the conceptualization and the development of the idea in the manuscript. Y.Z. performed the real data analysis. A.B., C.X., and K.E. provided data resources, biological interpretation, and funding support. All co-authors have reviewed the manuscript.

## Acknowledgments

H.D., K.E., Z.H. are supported by R01HL153042 and R01AR079220.

H.D., C.X., Z.H. are supported by FL DEPT OF HLTH BIOMED RES PGM/J&E KING 21K11.

Large Language Models (i.e., chatGPT) were used to correct written text (spell checkers, grammar checkers).

